# Structural changes following the reversal of a Y Chromosome to an autosome in *Drosophila pseudoobscura*

**DOI:** 10.1101/058412

**Authors:** Ching-Ho Chang, Amanda M. Larracuente

**Author notes:** Author for Correspondence: Department of Biology, University of Rochester, Rochester USA.

## Abstract

Robertsonian translocations resulting in fusions between sex chromosomes and autosomes shape karyotype evolution in animals by creating new sex chromosomes from autosomes. These translocations can also reverse sex chromosomes back into autosomes, which is especially intriguing given that autosomes and sex chromosomes differ in gene regulation and chromatin environment. While researchers are beginning to understand X chromosomes reversals to autosomes at a genomic level, it is difficult to study reversals of Y chromosomes because of their rapid sequence turnover and high repeat content. To gain insight into the genomic events following a Y chromosome reversal, we investigated an autosome-Y translocation in a well-studied and tractable organism, *Drosophila pseudoobscura*. About 10-15 Mya, the ancestral Y chromosome fused to a small autosome (the dot chromosome) in an ancestor of *D. pseudoobscura*. We used single molecule real-time sequencing reads to assemble the genic part of the *D. pseudoobscura* dot chromosome, including this Y-to-dot translocation. We find that the intervening sequence between the ancestral Y and the rest of the dot chromosome is only ~78 Kb and has a low repeat density, suggesting that the centromere now falls outside, rather than between, the fused chromosomes. The Y-to-dot region is 100 times smaller than the *D. melanogaster* Y chromosome, owing to repeat landscape changes. Previous studies suggest that recurrent selective sweeps favoring shorter introns helped to shrink the Y-to-dot following the translocation. Our results suggest that genetic drift and a small ancestral Y chromosome may also help explain the compact size of the Y-to-dot translocation.

## Introduction

Most sex chromosomes evolve from a pair of homologous autosomes, where one chromosome acquires a sex-determining locus, suppresses recombination with its homolog and eventually degenerates into a differentiated Y (or W in cases of female heterogamety) chromosome (reviewed in Ohno 1967; Charlesworth and Charlesworth 1978; Bull 1983; Charlesworth and Charlesworth 2000). While sex chromosomes appear stable in some taxa (*e.g*. mammals), they can be labile in others (*e.g*. non-mammalian vertebrates Schartl 2004; Ezaz, et al. 2006; insects White 1973; Vicoso and Bachtrog 2015). Robertsonian translocations between sex chromosomes and autosomes are important for sex chromosome evolution—the resulting chromosome fusions create new sex chromosomes from autosomes and new autosomes from sex chromosomes (Vicoso and Bachtrog 2013, 2015). X-autosome translocations are well-documented in some mammals (*e.g*. Indian muntjacs, Wurster and Benirschke 1970; shrews, Ford, et al. 1957; gerbils, Wahrman and Zahavi 1955; voles, Fredga 1970), sticklebacks (Ross, et al. 2009) and *Drosophila* species (White 1973; Steinemann 1982; McAllister and Charlesworth 1999; Vicoso and Bachtrog 2015), where autosome-sex chromosome fusions have created new sex chromosomes (i.e. neo-sex chromosomes) of independent origins and different ages.

Following the fusion of an autosome to a sex chromosome, the neo-Y chromosome usually degenerates and loses many of its genes relative to its counterpart, the neo-X chromosome. This degeneration occurs as a result of suppressed recombination that leads to a reduced efficacy of natural selection and the accumulation of deleterious mutations and repeats (transposable elements and satellite DNAs; reviewed in Charlesworth and Charlesworth 2000; Steinemann and Steinemann 2005; Bachtrog 2013). The degenerated Y chromosome is a harsh genomic environment for most genes—it is dense in repetitive elements, heterochromatic and nonrecombining (Bonaccorsi, et al. 1988; Bonaccorsi and Lohe 1991). Old differentiated Y chromosomes like *D. melanogaster*’s are gene poor (*e.g*. up to 20 genes in 40 Mb; Hoskins, et al. 2002). Only Y-linked genes that retain or evolve male-related functions are expected to survive on the Y chromosome; and these genes can have peculiar structures—some *Drosophila* Y-linked genes have introns that are Megabases long (hereon referred to as mega-introns) and filled with tandem repeats (Bonaccorsi, et al. 1988; Bonaccorsi, et al. 1990; Bonaccorsi and Lohe 1991; Kurek, et al. 1998; Kurek, et al. 2000; Reugels, et al. 2000; Piergentili 2007).

Sex chromosomes differ from major autosomes in gene regulation and ploidy (reviewed in Steinemann and Steinemann 2005) but despite these gross differences, sex chromosomes can revert back into autosomes. The small heterochromatic *Drosophila* autosome (dot chromosome; also known as Muller F) is an old X chromosome that reverted to an autosome in an ancestor of Drosophilids (Vicoso and Bachtrog 2013). In many ways, the dot chromosome still has more in common with an X chromosome than other autosomes (Larsson and Meller 2006; Riddle and Elgin 2006; Vicoso and Bachtrog 2013). While researchers are just beginning to understand the process of X chromosome reversal, Y chromosome reversals are relatively unexplored. *D. pseudoobscura* offers a unique opportunity to study Y chromosome reversals. Two chromosomal rearrangements occurred 10-15 Mya in an ancestor of *D. pseudoobscura:* an X-autosome and an autosome-Y translocation (Carvalho and Clark 2005; Larracuente, et al. 2010). The X-autosome translocation created a pair of neo-sex chromosomes: the current Y is not homologous to the ancestral Y chromosome (Carvalho and Clark 2005) and instead may be a neo-Y derived from the homolog of the autosome that fused to the X chromosome (Larracuente, et al. 2010). The ancestral Y chromosome translocated to the dot chromosome (Larracuente, et al. 2010). Thus after 60 Myr of paternal transmission, the ancestral Y chromosome reverted to an autosome and is now passed through both sexes. Because there are no known transition stages in any extant *obscura* group species, we do not know which event—the X-autosome or autosome-Y translocation—came first (Larracuente, et al. 2010). Following the autosome-Y translocation, the former Y chromosome (hereon referred to as the Y-to-dot region) may have shrunk 100-fold in size (Carvalho and Clark 2005; Larracuente, et al. 2010). Patterns of nucleotide variability in the Y-to-dot region are consistent with a history of selective sweeps, suggesting that positive selection may have favored large deletions of intronic sequences after becoming autosomal (Larracuente and Clark 2014). Therefore, *D. pseudoobscura* offers a window into the evolutionary events following a Y chromosome reversal.

Studying the structural changes that occurred after the Y-to-dot translocation has been difficult because this region is poorly assembled (Larracuente, et al. 2010). Heterochromatic sequences rich in repeats are often underrepresented in BAC libraries and cloning vectors (Brutlag, et al. 1977; Lohe and Brutlag 1986, 1987) used in traditional sequencing methods, making them difficult to assemble (Hoskins, et al. 2002). The short reads from next generation sequencing technologies have difficulty spanning and assembling repeats (Treangen and Salzberg 2012). Single molecule real-time sequencing technologies (SMRT) circumvent some of the problems with traditional and next generation short read sequencing (Eid, et al. 2009). SMRT reads from Pacific Biosciences (PacBio) are on average ~16 Kb long, but reach up to 50 Kb (with current technology). These long read lengths help span repeats and assemble repetitive regions (Schatz, et al. 2010; Chaisson, et al. 2014).

To infer the origins of the Y-to-dot translocation and detail the genomic changes accompanying this Y chromosome reversal, we used PacBio SMRT reads to assemble the entire genic portion of the *D. pseudoobscura* dot chromosome, including the Y-to-dot region. We reveal the breakpoints between the conserved part of the dot chromosome and the Y-to-dot region and discover that the region between them spans only ~78 Kb. Based on the organization of this region, we infer that at least one chromosomal inversion followed a Robertsonian translocation between the ancestral Y and dot chromosomes. We also show that while the Y-to-dot is 100-fold smaller than the *D. melanogaster* Y chromosome, the distribution of intron sizes does not differ from that of the Y chromosome, outside of the mega-introns.

## Results

### Assembly of dot-Y

We refer to the entire dot chromosome in species that have the translocation as the dot-Y chromosome. To determine the complete sequence of the dot-Y chromosome in *D. pseudoobscura*, we assembled the genome using ~70X coverage PacBio long reads. Our assembly contains 612 contigs covering 155.9Mb (N50 = 1.35 Mb, Table 1). All 5 ancestral Y-linked genes, all 79 conserved dot chromosome genes and telomeric TART transposons are found on a single 1.92 Mb contig in our assembly (contig 000025F). This single contig appears on 62 different scaffolds in the latest Flybase reference (r3.03; English, et al. 2012; Fig S1. and Table S1). We found all 5 Y-to-dot genes in a 320-Kb centromere-proximal region of the dot-Y chromosome (Fig 1). The most distal gene in our contig is *Plex-A*, consistent with a previous study (Villasante, et al. 2007). The orientation of these 5 Y-to-dot genes is also the same as previous reports (Larracuente, et al. 2010). Based on this assembly, we define the formerly Y-linked region from the start of the contig to *Ppr-Y* as Y-to-dot, and the conserved gene-dense region from *Cadps* to *Plex-A* as conserved-dot (Fig 1). Our assembly of the Y-to-dot and conserved-dot regions is consistent with our expectations with respect to gene content and order. However, the region containing the breakpoint between the two sections of the dot-Y revealed an interesting structure.

**Figure 1.**
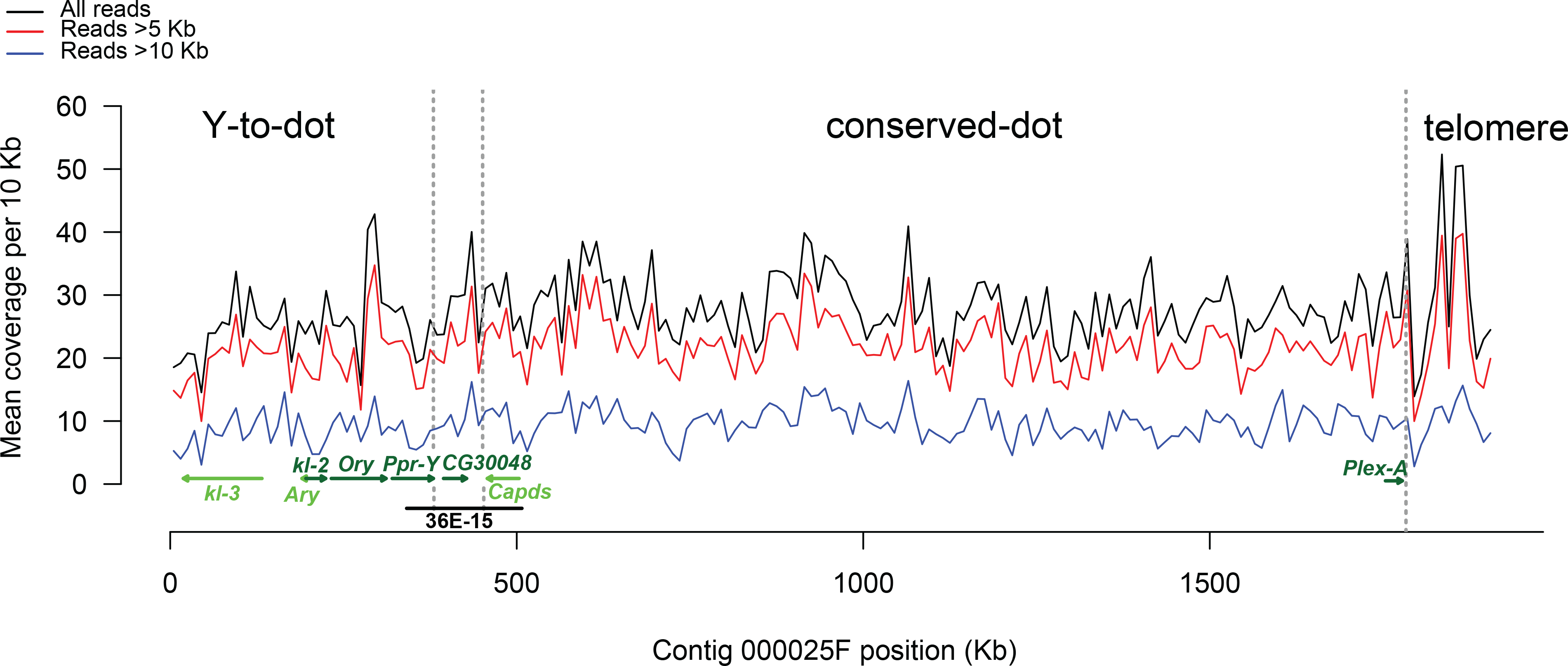
The structure and gene orientation of the dot-Y chromosome in *D. pseudoobscura*. The lines correspond to the coverage of all reads, reads >5 Kb, and reads > 10 Kb across the entire dot-Y chromosome. The Y-to-dot, conserved-dot and telomeric regions are delineated by gray dotted lines. All 5 Y-to-dot genes, the duplicated gene between the Y-to-dot and conserved dot, 2 conserved-dot genes (the most proximal and distal) and their orientations are indicated with green lines. The coordinates of the *D. persimilis* BAC (36E-15) are indicated with a black line.

**Table 1.**
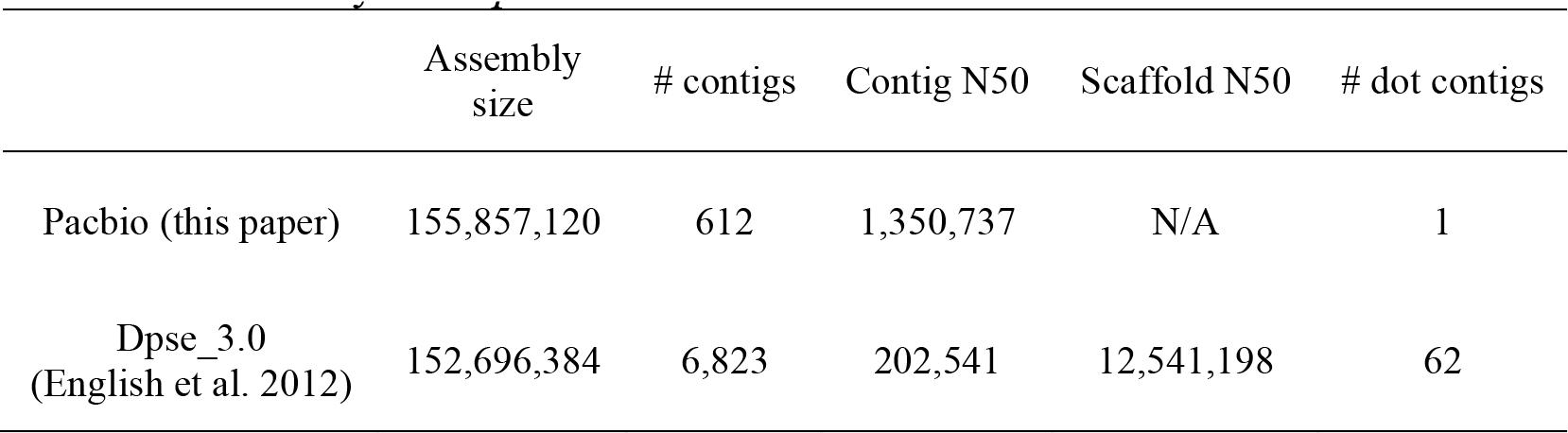
Summary of *D. pseudoobscura* assemblies

|  | Assembly size | # contigs | Contig N50 | Scaffold N50 | # dot contigs |
| --- | --- | --- | --- | --- | --- |
| Pacbio (this paper) | 155,857,120 | 612 | 1,350,737 | N/A | 1 |
| Dpse_3.0<br>(English et al. 2012) | 152,696,384 | 6,823 | 202,541 | 12,541,198 | 62 |

First, the region between the Y-to-dot and the conserved-dot is only ~78 Kb. This is surprising, given that we initially expected the centromere to fall between these two parts of the chromosome if it originated from a Robertsonian translocation. The region between the Y-to-dot and conserved-dot has two recent large duplications containing a 3,193-bp exon from a testis-specific gene—an ortholog of *CG30048* in *D. melanogaster* (the 3 fragments are 5,194 bp, 10,765 bp and 10,548 bp long and are > 95% identical; Fig S2). Interestingly, *CG30048* (TCONS_00043496 and TCONS_00043497 in our annotation) is a parental copy of an ancestral Y-linked gene—*Polycystein-related Y*, or *PRY*. In *D. pseudoobscura, PRY* is on chromosome *XL* and has testis-specific expression (Fig S3). No substitutions differentiate the two duplicates of *CG30048*. While one copy of *CG30048* is highly expressed in testes (FPKM=58 in testes but <3 in male carcass or females, Table S2), the other copy lacks the first 2 exons and is not significantly expressed (all transcriptomes FPKM<3, Table S2). These recent large duplications likely prevented the assembly of this region using *Illumina* and *Sanger* sequencing reads.

To confirm our assembly, we surveyed BACs from *D. pseudoobscura* and its sister species, *D. persimilis*. We found 4 BACs (CH226-45K5, CH226-45K6, CH226-6C6, CH226-55K10) and 3 fosmids (CH1226-51D24, CH1226-62C1, CH1226-33C10) from *D. pseudoobscura* that extend into region between the Y-to-dot and conserved dot (Table S3). However, we found that a single BAC from *D. persimilis* (TSC#14011-0111.49 0036E-15) spans the entire region and contains at least part of 3 dot-Y genes, including *Ppr-Y, Cadps* and *Dyrk3* (Fig 1). This BAC agrees with the structural arrangement of our assembly— the average insert size of this BAC library is 151 Kb (Song, et al. 2011) and according to our assembly, the PCR fragments from 3 genes spans 187 Kb in *D. pseudoobscura*. It also further supports that the region between the Y-to-dot and conserved-dot is conserved in *D. pseudoobscura* and its closely related species, *D. persimilis* (Larracuente, et al. 2010).

### Chromosomal rearrangement

The movement of Y-linked genes could have been the result of a large segmental duplication to the dot chromosome, a Robertsonian translocation or another chromosome fusion event. Because the organization of the Y chromosome appears to change rapidly over short periods of evolutionary time, it is difficult to infer the initial event. It is unlikely that a large segmental duplication moved the Y-linked genes to the dot chromosome because in *D. melanogaster*, these genes span both Y chromosome arms. Though the ancestral Y-linked genes are on the dot-Y, the rDNA and their intergenic spacer (IGS) sequences typically found on *Drosophila* Y chromosomes are absent from this region cytologically (Larracuente, et al. 2010) and in our assembly. Instead, the rDNA IGS repeats are on the current Y chromosome of *D. pseudoobscura*, suggesting that these sequences transferred to the Y chromosome either from the Y-to-dot or from the X chromosome (Larracuente, et al. 2010).

Directly following a Robertsonian translocation, the position of the centromere should fall between the Y-to-dot and conserved-dot regions. To ask if the region between the Y-to-dot and conserved-dot contains centromere-derived sequences, we determined the distribution of repetitive elements (satellite DNAs and transposons) across the dot-Y chromosome. Typical *Drosophila* centromeres are large (~400 Kb) and enriched in satellite DNA and transposable element-derived sequences (Karpen and Allshire 1997). The Y chromosome centromere in the *melanogaster* group is derived from telomeric retrotransposons (Abad, et al. 2004). However, the 78-Kb region between the Y-to-dot and conserved-dot is not enriched in repetitive sequences (Fig 2; Kruskal Wallis test and multiple comparison, Bonferroni *P* > 0.05)—we found no contiguous runs of tandem repeats longer than 5 Kb or TART/HeT-A arrays in this region (Table S4). Given its small size and relatively low density of repeats, we infer that the centromere is not located in the region between the Y-to-dot and conserved-dot.

**Figure 2.**
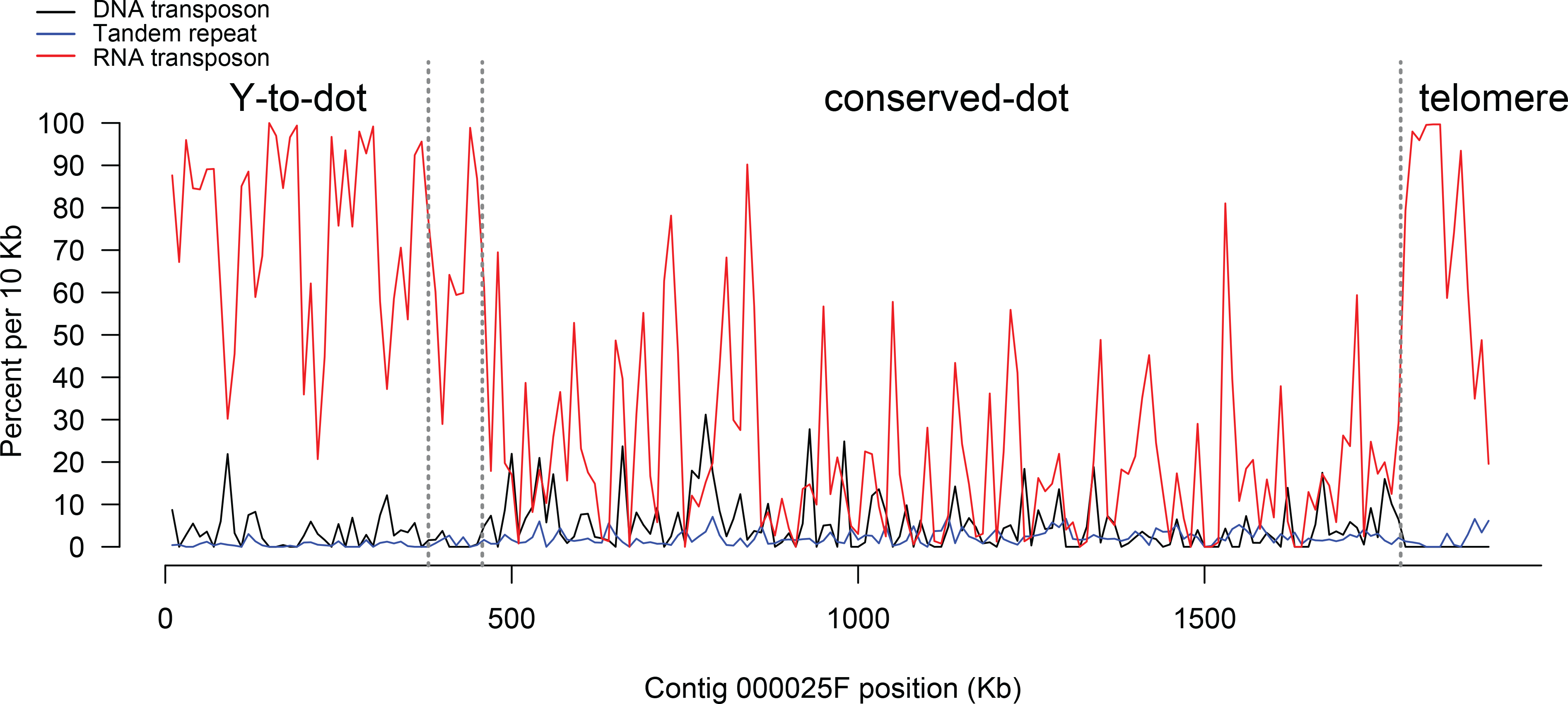
Repeat landscape across *D. pseudoobscura* dot-Y chromosome. The density of repeat types—DNA transposon, simple tandem repeats and RNA transposons—are plotted across the dot chromosome.

### Reorganization of the Y-to-dot region

The Y-to-dot region is 100-fold smaller than the *D. melanogaster* Y chromosome (Carvalho and Clark 2005). The 5 Y-to-dot genes span a total of ~320 Kb on the *D. pseudoobscura* dot chromosome, whereas their orthologs span both chromosome arms on the ~40 Mb *D. melanogaster* Y chromosome (Hoskins, et al. 2002). This drastic difference in size could be due to deletions in intergenic regions or deletions in both introns and intergenic regions following the translocation. We found a low density of tandem repeats in the Y-to-dot region (0.54% of total sequences; Fig 2), indicating that the deletion of repetitive DNA in intergenic regions contributes to the compact size of the Y-to-dot region. However, we surveyed the relative size of the Y and X chromosomes in karyotypes of *obscura* group species and find that some species have a small Y chromosome (Fig 3). Therefore, it is possible that the ancestral Y chromosome involved in the translocation was small.

**Figure 3.**
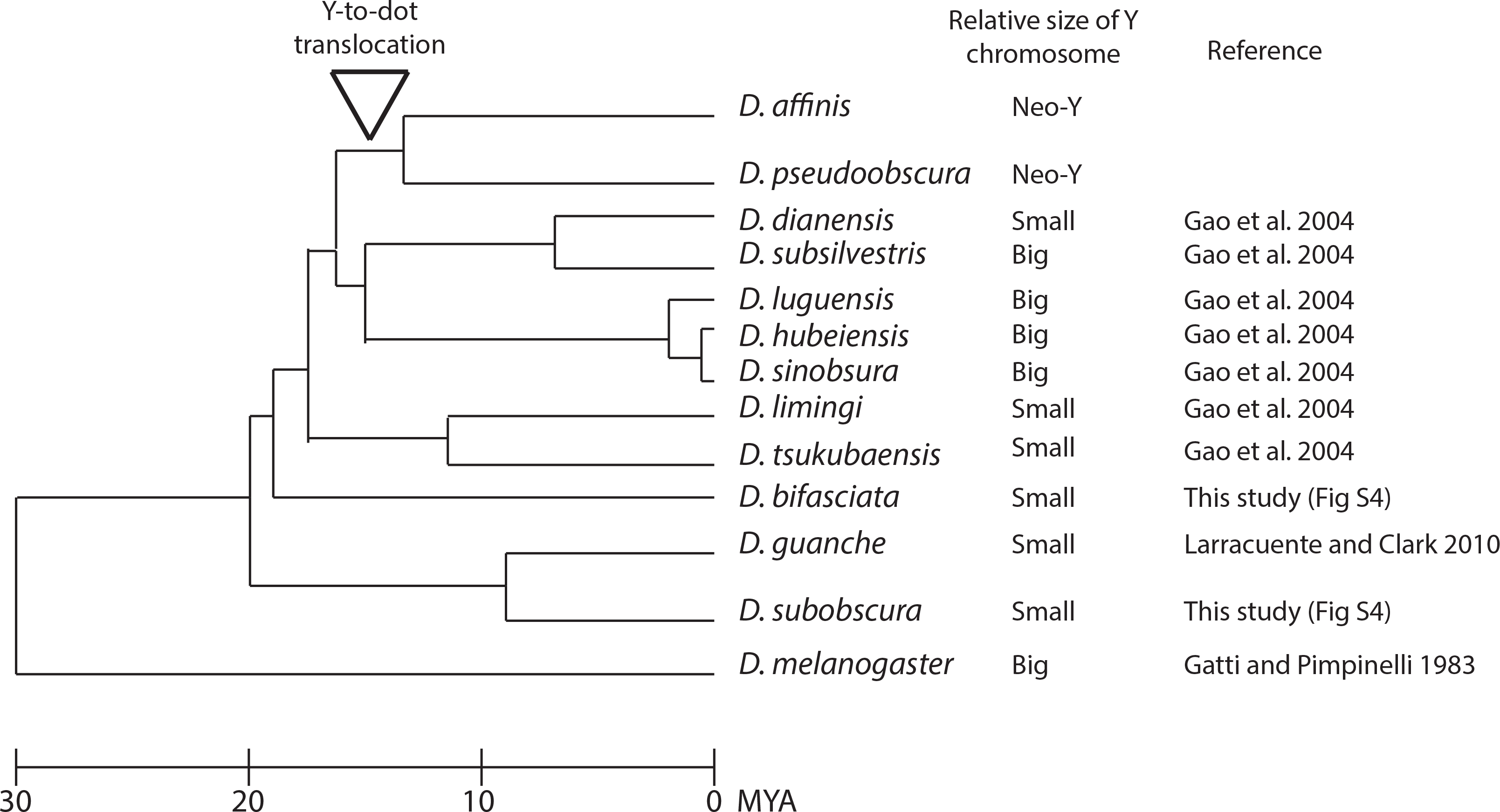
Evolution of Y chromosome size in the *obscura* group. The phylogeny is modified from (Gao, et al. 2007). The relative size of the ancestral Y chromosome compared to the ancestral X chromosomal arm (*i.e.* Muller A) in spreads of mitotic chromosomes (Fig S4; Gao, et al. 2004; Larracuente, et al. 2010). The ancestral *obscura* group Y chromosome may have been small.

Compared to intergenic regions, large introns should be under stronger selection due to costs of transcription (Prachumwat, et al. 2004). Some of the Y-to-dot genes appeared to shrink 10-fold after moving to the dot chromosome due to deletions in introns (Carvalho and Clark 2005; Larracuente, et al. 2010; Larracuente and Clark 2014). To quantify the contribution of intron size dynamics to this rearrangement, we studied intron sizes across the *D. pseudoobscura* dot-Y chromosome and compared the Y-to-dot genes to their orthologs on the *D. melanogaster* Y chromosome. As expected, introns across the dot-Y chromosome are larger than the other autosomes (median intron size dot-Y = 100.5 bp, autosomes = 68 bp; MWU, *P* = 4.726 ×10^-13^, Fig 4A), suggesting that the lack of crossing over on the dot contributes to larger introns. Within the *D. pseudoobscura* dot-Y chromosome, intron sizes differ among the Y-to-dot and conserved-dot regions (median intron size Y-to-dot = 1336 bp, conserved-dot = 93 bp; MWU, *P* = 4.449 ×10^-5^; Fig 4A). However, the *D. pseudoobscura* Y-to-dot genes do not have consistently smaller introns than Y-linked genes in *D. melanogaster* (Fig 3B and Table S5). While we fully assembled all 41 of the introns for the 5 Y-to-dot genes in *D. pseudoobscura*, only 22 of the 41 introns of their Y-linked orthologs in *D. melanogaster* (r6.03) are fully assembled. Of the 22 assembled Y-linked introns in *D. melanogaster*, 16 are larger in *D. pseudoobcura* (Fig 4B and Table S5). The introns of 3 Y-linked genes—*kl-3, kl-5* and *ORY (i.e. ks-1;* Kennison 1981)— contain Megabases of simple satellite sequences in *D. melanogaster* (Gatti and Pimpinelli 1983; Piergentili and Mencarelli 2008). Two of these mega-intron-containing genes, *kl-3* and ORY, are located in the *D. pseudoobscura* Y-to-dot region and are significantly smaller (~113 and 66 Kb, respectively). In our assembly, the largest intron in these genes is 41 Kb and no introns contain stretches of satellite DNA sequences. Therefore eliminating large introns, including the mega-introns, greatly affected the genome structure of Y-to-dot.

**Figure 4.**
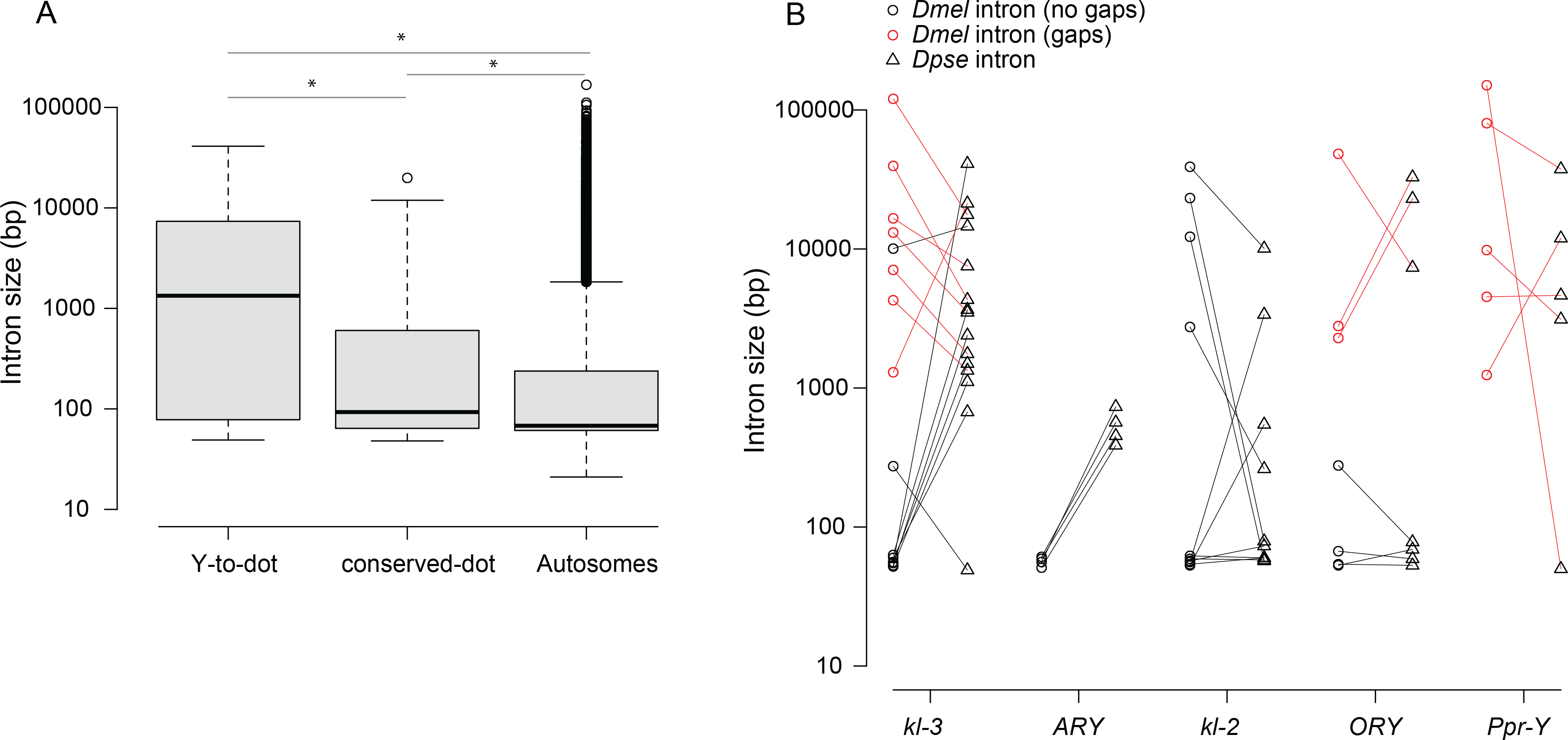
Intron size comparisons. A) Boxplots show intron size differences across the *D. pseudoobscura* genome. Intron sizes for the dot-Y chromosome regions are based on our PacBio assembly and annotations; autosomal introns are based on the r3.03 reference genome assembly and annotations (Flybase). Only introns from the first isoforms of each gene were used. For all dot-Y genes, we only plot those with orthologs in *D. melanogaster* (Blastx; e < 0.01). Asterisks (*) indicate significantly different means (Kruskal-Wallis; pair-wise Bonferroni; *P* < 0.05). B) Comparisons of individual orthologous intron pairs between the *D. melanogaster* Y chromosome (circles) and the *D. pseudoobscura* Y-to-dot translocation (triangles). Introns in the *D. melanogaster* r6.03 reference Y chromosome with gaps (indicated by Ns; and therefore are minimum intron sizes) are in red and those without gaps are in black. Orthologous introns between the species pair are connected with a line.

Though the intergenic and intronic regions of Y-linked genes are vastly different between *D. pseudoobscura* and *D. melanogaster*, the coding regions are conserved between species (Singh, et al. 2014). However, there is gene content turnover on *Drosophila* Y chromosomes (Koerich, et al. 2008), so the Y-to-dot translocation may have uncharacterized formerly Y-linked genes. To ask if there are new genes or non-coding RNAs associated with the translocation, we studied gene expression across the dot chromosome. The 5 Y-to-dot genes remained testis-specific after becoming autosomal, as previous studies suggested (Carvalho and Clark 2005). We found some evidence for one new gene in the Y-to-dot region—a predicted transcript originating from a duplication of CG9065 (TCONS_00020811) in the intron of *kl-3*. This duplication occurred after the Y-to-dot translocation because it shares 96% identity with its parental copy. However, this duplicate is not expressed (all RNA-seq datasets FPKM < 1, Table S2). Other than the original 5 Y-linked genes, no predicted genes in the translocated region show significant expression (FPKM>5; Table S2).

### Population genetic analyses

We estimated levels of nucleotide diversity and population recombination rates across the dot-Y chromosome for 11 strains of *D. pseudoobscura* and 12 strains of *D. miranda* (both species have a dot-Y). Due to a severe population bottleneck in the demographic history of *D. miranda* (Bachtrog and Andolfatto 2006; Jensen and Bachtrog 2011), nucleotide diversity is lower than in *D. pseudoobscura*. In both species, the major autosomes (Muller B and E) harbor more nucleotide diversity than the X (Muller A and D) and dot-Y chromosomes (median *π_non-dot autosomes_* = 0.0102 and 0.0014 in *D. pseudoobscura* and *D. miranda*, respectively; median *π_X_*= 0.0081 and 0.0012 in *D. pseudoobscura* and *D. miranda*, respectively; median *π_dot-Y_* = 0.0008 and 0.0002 in *D. pseudoobscura* and *D. miranda*, respectively; MWU, *P* < 2.2 ×10^−16^). Within the dot-Y chromosome, nucleotide diversity is ~2-fold greater on the conserved-dot compared to the Y-to-dot (*π_conserved-dot_* = 0.0009, *π_Y-to-dot_* = 0.0005 in *D. pseudoobscura*; *π_conserved-dot_* = 0.0002, *π_Y-to-dot_* = 0.0001 in *D. miranda;* Kruskal Wallis test; multiple comparisons, Bonferroni *P* > 0.05, Fig 5). Consistent with previous reports (Larracuente and Clark 2014), we detect a low recombination rate on the dot chromosomes of both *D. pseudoobscura* (Rmin=152) and *D. miranda* (Rmin=17) along the 1.9 Mb dot chromosome. The low recombination rates likely contribute to the reduced nucleotide diversity across these dot chromosomes.

**Figure 5.**
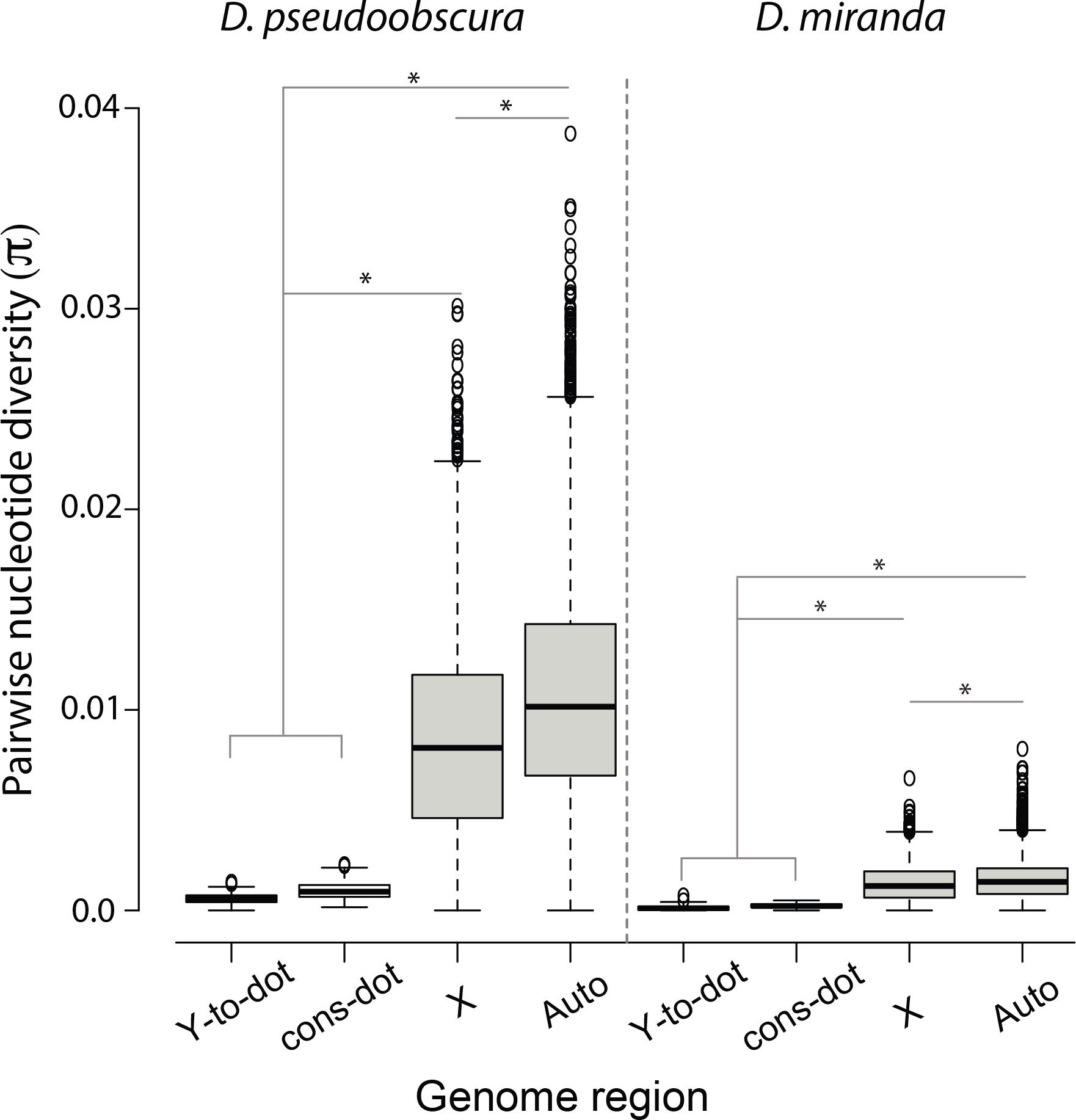
Nucleotide diversity across the genome in *D. pseudoobscura* and *D. miranda*. The dot chromosome, including Y-to-dot and conserved-dot regions, have lower pairwise nucleotide diversity than representative autosomes (those chromosome arms not involved in sex-autosome translocation; *i.e.* Muller B and E) and the X chromosome *(i.e.* Muller A and D). Asterisks (*) indicate significantly different means (Kruskal-Wallis; pair-wise Bonferroni; *P* < 0.05).

## Discussion

Robertsonian translocations occur through the fusion of two acrocentric or telocentric chromosomes to create one metacentric chromosome. If the Y-to-dot event was a Robertsonian translocation, then the fused centromeres of the Y and dot chromosomes would have resided between the two chromosomal arms immediately after the translocation occurred. However, we found that the region between the Y-to-dot and conserved-dot is relatively small, lacks highly repetitive DNA and therefore is unlikely to correspond to the centromere. A pericentric inversion may have moved the centromere to other side of the translocated region, or the centromere itself may have shifted positions. Without knowing the gene order on the ancestral Y chromosome involved in the translocation, it will be hard to determine definitively—the event is 10-15 Myr old, leaving time for subsequent rearrangements. Alternatively, the dot and Y chromosomes may have fused as a result of an unequal crossover event between the telomeric region of the Y chromosome and the centromeric region of the dot chromosome. While we do not know the identity of the current dot chromosome centromere in *D. pseudoobscura*, the latter hypothesis would predict that it is derived from the ancestral Y chromosome centromere. Further studies in the *obscura* group are necessary to reveal the detailed evolutionary history of this event.

Sex-autosome translocations are often deleterious mutations that do not persist in populations (Ohno 1967). The Y-to-dot translocation therefore seems like a relatively rare event in that it was successful and fixed in the ancestral population. For a male-specific gene, the Y chromosome offers shelter from relaxed or antagonistic selection in females (Fisher 1931; Rice 1987) but at the cost of a reduced efficacy of natural selection (Charlesworth and Charlesworth 2000). Autosomes offer larger effective population sizes and higher recombination rates that should facilitate more efficacious selection (Betancourt and Presgraves 2002; Presgraves 2005). However, these benefits are only realized over long evolutionary timescales, not immediately following a chromosome translocation. What then made the Y-to-dot translocation successful? The Y-to-dot does not represent the only gene traffic between the dot and Y chromosome in *Drosophila* species. There are at least three independent cases of individual gene movements between the dot and Y chromosomes: *kl-5* (Dyer, et al. 2011), *JYalpha* (Carvalho and Clark 2013) and *PRY* (Leung, et al. 2015). The dot chromosome may offer a suitable environment for a sex-linked gene. The dot and X chromosomes have similarities: they both have chromosome-specific gene regulation, are haplosufficient, and feminizing in intersexes (adding an X or dot to a fly with 2X:3A biases development toward females; reviewed in Larsson and Meller 2006;

Riddle and Elgin 2006; Vicoso and Bachtrog 2013). The dot may also have some similarities to Y chromosomes in chromatin environment—they are both largely heterochromatic (80% of the dot chromosome is heterochromatic) and, at least in some species, they share satellite DNAs (*e.g*. Lohe, et al. 1993). A heterochromatic environment is required for the proper regulation of some autosomal genes (*e.g*. Eberl, et al. 1993), and Y-linked genes may have similar requirements. It is possible that the initial dot-Y fusion was successful because it placed the Y-to-dot region in a heterochromatic environment similar enough to the ancestral Y chromosome. While the Y and dot chromosomes may be similar in their heterochromatic regions, the Y-linked genes are now located relatively close to the conserved-dot region. In *D. melanogaster*, the gene-dense region of the dot is a unique type of chromatin with properties of both heterochromatin and euchromatin (Sun, et al. 2000; Leung, et al. 2010). Following the translocation, the formerly Y-linked genes retained testis specific expression but their structure changed dramatically: some individual genes are at least 10-fold smaller than their orthologs on the *D. melanogaster* Y chromosome due to deletions in very large introns. The shortening of large introns following the Y-to-dot translocation may have been driven by recurrent selective sweeps (Larracuente and Clark 2014).

Introns across the genome are expected to evolve under natural selection—many contain regulatory sequences and are constrained by splicing requirements and transcription costs (Prachumwat, et al. 2004). As such, there is a genome-wide correlation between intron size and recombination rate in *Drosophila* (Carvalho and Clark 1999; Comeron and Kreitman 2000; also see Comeron and Kreitman 2002). This may be especially true for highly expressed genes (Castillo-Davis, et al. 2002; Urrutia and Hurst 2003). Consistent with this observation, we found that intron sizes are larger on the dot chromosome—where meiotic recombination *via* crossing over is at most very low—compared to other autosomes. The mega-introns of *Drosophila* Y-linked genes may be consequences of the reduced efficacy of purifying selection (Carvalho and Clark 1999; Carvalho 2003). Alternatively, these mega-introns may operate under different selection pressures, as some have hypothesized that they serve a role in spermatogenesis (Bonaccorsi, et al. 1990; Pisano, et al. 1993; Piergentili, et al. 2004). Our analysis lends some general insights into the selection pressures on intron sizes. After becoming autosomal for 10-15 My, outside of lacking the few Y-linked mega-introns and very large introns, most introns in the Y-to-dot region are not smaller than their Y-linked orthologs. In many cases, the introns are larger in the Y-to-dot region—consistent with the earlier observation that the correlation between recombination rate and intron size does not hold for large introns (Carvalho and Clark 1999). This suggests that in *D. pseudoobscura*, purifying selection against large introns (*e.g*. below ~10 Kb) on the Y-to-dot is inefficient. *D. pseudoobscura* presumably has mega-introns—large lampbrush-like loops are formed during spermatogenesis (Piergentili 2007)—but we do not find evidence that they come from the homologous introns in the Y-to-dot regions. Lampbrush-like loops originate from different Y-linked introns in *D. melanogaster* and *D. hydei* (Reugels et al. 2000). This suggests that *D. pseudoobscura* acquired these loops independently, perhaps on the new Y chromosome.

Whereas the Y-to-dot genes appeared to shrink 10-fold, the entire Y-to-dot region appeared to shrink ~100-fold (compared to the *D. melanogaster* Y chromosome). This suggests that the Y-to-dot intergenic regions were severely reduced after becoming autosomal. However, our view of *Drosophila* Y chromosomes is *melanogaster-centric*. The ancestral *obscura* Y chromosome could have been structurally different from the *D. melanogaster* Y chromosome. It is possible that the compact size of the Y-to-dot region is not only due to intergenic deletions of a large ancestral Y chromosome, but instead that the ancestral *obscura* group Y chromosome that fused to the dot chromosome was itself small. Y chromosome size varies among *Drosophila* species and even within species (White 1973). The karyotypes of other *Drosophila* species are consistent with the hypothesis that the ancestral *obscura* group Y chromosome was small (Fig 3 and Fig S4). However, without more genomic information from these Y chromosomes, it will be difficult to infer their evolutionary dynamics. Therefore, a combination of selection favoring deletions in large introns, genetic drift and perhaps even a small ancestral Y chromosome may explain the structural differences between the Y-to-dot region and the *D. melanogaster* Y chromosome.

## Materials and methods

### PacBio assemblies

We downloaded raw PacBio reads from ftp://ftp.hgsc.bcm.edu/Dpseudoobscura/Towards_finishing_the_D.pseudoobscura_genome/PacBio_Data/FastQfiles/ (data generated in 2014; permission to use these reads was generously granted by Drs. Steve Schaeffer and Stephan Richards; Table S6). This data set is 10.8 Gb of sequence in total and includes 1.6 million reads with read lengths averaging 6.7±4.2-Kb long (excluding reads < 500 bp). We used the Falcon assembler v0.3.0 (https://github.com/PacificBiosciences/FALCON-integrate) to filter, correct and assemble reads (FALCON configuration file in Supplemental Doc 1). We polished the assembly using Quiver (SMRT Analysis v2.3.0; Chin, et al. 2013). To compare our PacBio assembly with the latest *D. pseudoobscura* genome assembly (r3.03; English, et al. 2012; Flybase), we used MUMMER (v3.23) nucmer with the parameters “-1200 -D 20 --maxmatch -nosimplify” (Kurtz, et al. 2004). We also annotated repetitive DNA using RepeatMasker 4.06 with Repbase 20150807 and the parameters “-species drosophila -s -pa 10” (Smit, et al. 2013).

### Annotation andRNA-seq analysis

We identified exon-introns junctions by mapping annotated transcripts (r3.03) and transcripts of Y-linked genes from NCBI (gi|295126506|gb|GU937420.1|, gi|255764727|gb|EU595397.2|, gi|295126512|gb|GU937423.1|) to our assembly using exonerate 2.4.0 (Slater and Birney 2005). To complement these data, we mapped RNA-seq reads (Chen, et al. 2014; Table S6) to our assembly using Tophat v2.0.13 with the parameters “-p 16 -b2-very-sensitive” (Trapnell, et al. 2009). We annotated transcripts, eliminated small redundant isoforms (gffread parameters “-M-K”) and estimated expression level with cufflinks v2.2.2 (parameters “-p 16 -u -b”) (Trapnell, et al. 2012). We manually modified gene annotations in the Y-to-dot region due to their longer introns. We then searched for homologous genes using the predicted transcripts as BLAST queries to all *D. melanogaster* translations and transposon sequences (r6.07) with NCBIblast (v2.2.30+; (Altschul, et al. 1990)). To survey BAC sequences, the *Ppr-Y* primers were obtained from (Larracuente and Clark 2014), and the *Cadps* and *Dyrk3* primers are listed in Table S7. Our dot chromosome assembly and gtf file containing annotations are available on our website (http://blogs.rochester.edu/larracuente/lab-resources-and-links) and in the Dryad Digital Repository.

### Population analyses and phylogenetic analysis

We downloaded population genomic datasets including 11 *D. pseudoobscura* individuals from 7 different populations (McGaugh and Noor 2012; Zhou and Bachtrog 2012; Smukowski Heil, et al. 2015; NCBI accession numbers in Table S6). After trimming with Trim Galore (v0.4.0; http://www.bioinformatics.babraham.ac.uk/projects/trim_galore/), we mapped all reads to the genome using a pipeline modified from (Lack, et al. 2015; https://github.com/justin-lack/Drosophila-Genome-Nexus/blob/master/round1_assembly.pl). Briefly, we used bwa mem (v0.7.12; Li and Durbin 2010), picard (v1.114; http://picard.sourceforge.net.) and stampy (v1.0.23; Lunter and Goodson 2011). To improve the sensitivity, bam files from the same species were used to call SNPs with GATK UnifiedGenotyper with multi-sample parameters “-mbq 10 - stand_call_conf 31 -stand_emit_conf 31 -ploidy 2” (v3.4-46; McKenna, et al. 2010). We extracted SNP information from vcf files using Vcftools (v0.1.14; Danecek, et al. 2011) and calculated Rmin and pairwise nucleotide diversity (n) in 10 Kb windows using compute 0.8.4 (Thornton 2003). We performed statistical analyses and plotted figures in R (v3.1.2) with ‘agricolae’ (v1.2-3; https://cran.r-project.org/web/packages/agricolae/index.html) and ‘ggplot2’ (v2.1.0).

We either downloaded *PRY, CG30048* and *CG33482* and their orthologs from 10 *Drosophila* species from Flybase (FB2015_04) or extracted them from genome assemblies (this study). We downloaded the *Ceratitis capitata* (L0C101450228) and *Musca domestica* sequences (MD0A003316, Scott, et al. 2014) from NCBI and Vectorbase, respectively. We used Mega (v6.06; Tamura, et al. 2013) to align amino acid sequences manually, and construct a phylogeny by Maximum likelihood with JTT model and 1000 bootstraps.

### Karyotyping

We performed brain squashes according to Larracuente and Ferree (2015) with some modifications. We dissected 3~5 male third instar larvae, and transferred brains to a hypotonic solution of 0.5% sodium citrate for 8~10 min. We fixed the brains in 45% acetic acid for 10~20 min and then squashed, snap froze in liquid nitrogen and dehydrated the slides in absolute ethanol for more than 30 min. Slides were mounted in VectaShield with DAPI (Vector Laboratories), visualized on a Leica DM5500 fluorescence microscope at 100X, imaged with a Hamamatsu Orca R2 CCD camera and analyzed using Leica’s LAX software.

## Acknowledgements

We thank Drs. Stephan Richards and Steve Schaeffer for sharing the *D. pseudoobscura* PacBio reads—these data were generated under the support of a National Institutes of Health National Institute of General Medical Sciences grant R01 GM098478 to S.W. Schaeffer. We also thank the University of Rochester Center for Integrated Research Computing for access to computing cluster resources.

## References

Abad JP, de Pablos B, Agudo M, Molina I, Giovinazzo G, Martin-Gallardo A, Villasante A. 2004. Genomic and cytological analysis of the Y chromosome of Drosophila melanogaster: telomere-derived sequences at internal regions. Chromosoma 113:295–304.

Altschul SF, Gish W, Miller W, Myers EW, Lipman DJ. 1990. Basic local alignment search tool. J Mol Biol 215:403–410.

Bachtrog D. 2013. Y-chromosome evolution: emerging insights into processes of Y-chromosome degeneration. Nat Rev Genet 14:113–124.

Bachtrog D, Andolfatto P. 2006. Selection, recombination and demographic history in Drosophila miranda. Genetics 174:2045–2059.

Betancourt AJ, Presgraves DC. 2002. Linkage limits the power of natural selection in Drosophila. Proceedings of the National Academy of Sciences of the United States of America 99:13616–13620.

Bonaccorsi S, Gatti M, Pisano C, Lohe A. 1990. Transcription of a satellite DNA on two Y chromosome loops of Drosophila melanogaster. Chromosoma 99:260–266.

Bonaccorsi S, Lohe A. 1991. Fine mapping of satellite DNA sequences along the Y chromosome of Drosophila melanogaster: relationships between satellite sequences and fertility factors. Genetics 129:177–189.

Bonaccorsi S, Pisano C, Puoti F, Gatti M. 1988. Y chromosome loops in Drosophila melanogaster. Genetics 120:1015–1034.

Brutlag D, Carlson M, Fry K, Hsieh TS. 1977. DNA-Sequence Organization in Drosophila Heterochromatin. Cold Spring Harbor Symposia on Quantitative Biology 42:1137–1146.

Bull JJ. 1983. Evolution of sex determining mechanisms. Menlo Park, CA: The Benjamin/Cummings Publishing Company, INC.

Carvalho AB. 2003. The advantages of recombination. Nat Genet 34:128–129.

Carvalho AB, Clark AG. 2013. Efficient identification of Y chromosome sequences in the human and Drosophila genomes. Genome Res 23:1894–1907.

Carvalho AB, Clark AG. 1999. Intron size and natural selection. Nature 401:344.

Carvalho AB, Clark AG. 2005. Y chromosome of D. pseudoobscura is not homologous to the ancestral Drosophila Y. Science 307:108–110.

Castillo-Davis CI, Mekhedov SL, Hartl DL, Koonin EV, Kondrashov FA. 2002. Selection for short introns in highly expressed genes. Nat Genet 31:415–418.

Chaisson MJ, Huddleston J, Dennis MY, Sudmant PH, Malig M, Hormozdiari F, Antonacci F, Surti U, Sandstrom R, Boitano M, et al. 2014. Resolving the complexity of the human genome using single-molecule sequencing. Nature.

Charlesworth B, Charlesworth D. 2000. The degeneration of Y chromosomes. Philos Trans R Soc Lond B Biol Sci 355:1563–1572.

Charlesworth B, Charlesworth D. 1978. A model for the evolution of dioecy and gynodioecy. American Naturalist 112:975–997.

Chen ZX, Sturgill D, Qu J, Jiang H, Park S, Boley N, Suzuki AM, Fletcher AR, Plachetzki DC, FitzGerald PC, et al. 2014. Comparative validation of the D. melanogaster modENCODE transcriptome annotation. Genome Res 24:1209–1223.

Chin CS, Alexander DH, Marks P, Klammer AA, Drake J, Heiner C, Clum A, Copeland A, Huddleston J, Eichler EE, et al. 2013. Nonhybrid, finished microbial genome assemblies from long-read SMRT sequencing data. Nat Methods 10:563–569.

Comeron JM, Kreitman M. 2000. The correlation between intron length and recombination in Drosophila. Dynamic equilibrium between mutational and selective forces. Genetics 156:11751190.

Comeron JM, Kreitman M. 2002. Population, evolutionary and genomic consequences of interference selection. Genetics 161:389–410.

Danecek P, Auton A, Abecasis G, Albers CA, Banks E, DePristo MA, Handsaker RE, Lunter G, Marth GT, Sherry ST, et al. 2011. The variant call format and VCFtools. Bioinformatics 27:2156–2158.

Dyer KA, White BE, Bray MJ, Pique DG, Betancourt AJ. 2011. Molecular evolution of a Y chromosome to autosome gene duplication in Drosophila. Mol Biol Evol 28:1293–1306.

Eberl DF, Duyf BJ, Hilliker AJ. 1993. The role of heterochromatin in the expression of a heterochromatic gene, the rolled locus of Drosophila melanogaster. Genetics 134:277–292.

Eid J, Fehr A, Gray J, Luong K, Lyle J, Otto G, Peluso P, Rank D, Baybayan P, Bettman B, et al. 2009. Real-time DNA sequencing from single polymerase molecules. Science 323:133–138.

English AC, Richards S, Han Y, Wang M, Vee V, Qu J, Qin X, Muzny DM, Reid JG, Worley KC, et al. 2012. Mind the gap: upgrading genomes with Pacific Biosciences RS long-read sequencing technology. PLoS One 7:e47768.

Ezaz T, Stiglec R, Veyrunes F, Marshall Graves JA. 2006. Relationships between vertebrate ZW and XY sex chromosome systems. Curr Biol 16:R736–743.

Fisher RA. 1931. The evolution of dominance. Biol Rev 6:345–368.

Ford CE, Hamerton JL, Sharman GB. 1957. Chromosome polymorphism in the common shrew. Nature 180:392–393.

Fredga K. 1970. Unusual Sex Chromosome Inheritance in Mammals. Philosophical Transactions of the Royal Society B: Biological Sciences 259:15–36.

Gao J-j, Watabe H-a, Zhang Y-p, Aotsuka T. 2004. Karyotype differentiation in newly discovered members of the Drosophila obscura species group from Yunnan, China. Zoological Research 25:236–241.

Gao JJ, Watabe HA, Aotsuka T, Pang JF, Zhang YP. 2007. Molecular phylogeny of the Drosophila obscura species group, with emphasis on the Old World species. BMC Evol Biol 7:87.

Gatti M, Pimpinelli S. 1983. Cytological and genetic analysis of the Y chromosome of Drosophila melanogaster. I. Organization of the fertility factors. Chromosoma 88:349–373.

Hoskins RA, Smith CD, Carlson JW, Carvalho AB, Halpern A, Kaminker JS, Kennedy C, Mungall CJ, Sullivan BA, Sutton GG, et al. 2002. Heterochromatic sequences in a Drosophila whole-genome shotgun assembly. Genome biology 3:RESEARCH0085.

Jensen JD, Bachtrog D. 2011. Characterizing the influence of effective population size on the rate of adaptation: Gillespie’s Darwin domain. Genome Biol Evol 3:687–701.

Karpen GH, Allshire RC. 1997. The case for epigenetic effects on centromere identity and function. Trends Genet 13:489–496.

Kennison JA. 1981. The Genetic and Cytological Organization of the Y Chromosome of D. melanogaster. Genetics 98:529–548.

Koerich LB, Wang X, Clark AG, Carvalho AB. 2008. Low conservation of gene content in the Drosophila Y chromosome. Nature 456:949–951.

Kurek R, Reugels AM, Glatzer KH, Bunemann H. 1998. The Y chromosomal fertility factor Threads in Drosophila hydei harbors a functional gene encoding an axonemal dynein beta heavy chain protein. Genetics 149:1363–1376.

Kurek R, Reugels AM, Lammermann U, Bunemann H. 2000. Molecular aspects of intron evolution in dynein encoding mega-genes on the heterochromatic Y chromosome of Drosophila sp. Genetica 109:113–123.

Kurtz S, Phillippy A, Delcher AL, Smoot M, Shumway M, Antonescu C, Salzberg SL. 2004. Versatile and open software for comparing large genomes. Genome Biol 5:R12.

Lack JB, Cardeno CM, Crepeau MW, Taylor W, Corbett-Detig RB, Stevens KA, Langley CH, Pool JE. 2015. The Drosophila Genome Nexus: A Population Genomic Resource of 623 Drosophila melanogaster Genomes, Including 197 from a Single Ancestral Range Population. Genetics 199:1229–1241.

Larracuente AM, Clark AG. 2014. Recent selection on the Y-to-dot translocation in Drosophila pseudoobscura. Mol Biol Evol 31:846–856.

Larracuente AM, Ferree PM. 2015. Simple method for fluorescence DNA in situ hybridization to squashed chromosomes. JoVE 95:e52288.

Larracuente AM, Noor MA, Clark AG. 2010. Translocation of Y-linked genes to the dot chromosome in Drosophila pseudoobscura. Molecular Biology and Evolution 27:1612–1620.

Larsson J, Meller VH. 2006. Dosage compensation, the origin and the afterlife of sex chromosomes. Chromosome Res 14:417–431.

Leung W, Shaffer CD, Cordonnier T, Wong J, Itano MS, Slawson Tempel EE, Kellmann E, Desruisseau DM, Cain C, Carrasquillo R, et al. 2010. Evolution of a distinct genomic domain in Drosophila: comparative analysis of the dot chromosome in Drosophila melanogaster and Drosophila virilis. Genetics 185:1519–1534.

Leung W, Shaffer CD, Reed LK, Smith ST, Barshop W, Dirkes W, Dothager M, Lee P, Wong J, Xiong D, et al. 2015. Drosophila muller f elements maintain a distinct set of genomic properties over 40 million years of evolution. G3 (Bethesda) 5:719–740.

Li H, Durbin R. 2010. Fast and accurate long-read alignment with Burrows-Wheeler transform. Bioinformatics 26:589–595.

Lohe AR, Brutlag DL. 1987. Adjacent satellite DNA segments in Drosophila structure of junctions. J Mol Biol 194:171–179.

Lohe AR, Brutlag DL. 1986. Multiplicity of satellite DNA sequences in Drosophila melanogaster. Proc Natl Acad Sci U S A 83:696–700.

Lohe AR, Hilliker AJ, Roberts PA. 1993. Mapping simple repeated DNA sequences in heterochromatin of Drosophila melanogaster. Genetics 134:1149–1174.

Lunter G, Goodson M. 2011. Stampy: a statistical algorithm for sensitive and fast mapping of Illumina sequence reads. Genome Res 21:936–939.

McAllister BF, Charlesworth B. 1999. Reduced sequence variability on the Neo-Y chromosome of Drosophila americana americana. Genetics 153:221–233.

McGaugh SE, Noor MA. 2012. Genomic impacts of chromosomal inversions in parapatric Drosophila species. Philos Trans R Soc Lond B Biol Sci 367:422–429.

McKenna A, Hanna M, Banks E, Sivachenko A, Cibulskis K, Kernytsky A, Garimella K, Altshuler D, Gabriel S, Daly M, et al. 2010. The Genome Analysis Toolkit: a MapReduce framework for analyzing next-generation DNA sequencing data. Genome Res 20:1297–1303.

Ohno S. 1967. Sex chromosomes and sex-linked genes. Berlin, Heidelberg, New York: Springer-Verlag.

Piergentili R. 2007. Evolutionary conservation of lampbrush-like loops in drosophilids. BMC Cell Biol 8:35.

Piergentili R, Bonaccorsi S, Raffa GD, Pisano C, Hackstein JH, Mencarelli C. 2004. Autosomal control of the Y-chromosome kl-3 loop of Drosophila melanogaster. Chromosoma 113:188–196.

Piergentili R, Mencarelli C. 2008. Drosophila melanogaster kl-3 and kl-5 Y-loops harbor triple-stranded nucleic acids. J Cell Sci 121:1605–1612.

Pisano C, Bonaccorsi S, Gatti M. 1993. The kl-3 loop of the Y chromosome of Drosophila melanogaster binds a tektin-like protein. Genetics 133:569–579.

Prachumwat A, DeVincentis L, Palopoli MF. 2004. Intron size correlates positively with recombination rate in Caenorhabditis elegans. Genetics 166:1585–1590.

Presgraves DC. 2005. Recombination enhances protein adaptation in Drosophila melanogaster. Current biology : CB 15:1651–1656.

Reugels AM, Kurek R, Lammermann U, Bunemann H. 2000. Mega-introns in the dynein gene DhDhc7(Y) on the heterochromatic Y chromosome give rise to the giant threads loops in primary spermatocytes of Drosophila hydei. Genetics 154:759–769.

Rice WR. 1987. The accumulation of sexually antagonistic genes as a selective agent promoting the evolution of reduced recombination between primitive sex chromosomes. Evolution 41:911914.

Riddle NC, Elgin SC. 2006. The dot chromosome of Drosophila: insights into chromatin states and their change over evolutionary time. Chromosome Res 14:405–416.

Ross JA, Urton JR, Boland J, Shapiro MD, Peichel CL. 2009. Turnover of sex chromosomes in the stickleback fishes (gasterosteidae). PLoS Genet 5:e1000391.

Schartl M. 2004. Sex chromosome evolution in non-mammalian vertebrates. Curr Opin Genet Dev 14:634–641.

Schatz MC, Delcher AL, Salzberg SL. 2010. Assembly of large genomes using second-generation sequencing. Genome Res 20:1165–1173.

Scott JG, Warren WC, Beukeboom LW, Bopp D, Clark AG, Giers SD, Hediger M, Jones AK, Kasai S, Leichter CA, et al. 2014. Genome of the house fly, Musca domestica L., a global vector of diseases with adaptations to a septic environment. Genome Biol 15:466.

Singh ND, Koerich LB, Carvalho AB, Clark AG. 2014. Positive and purifying selection on the Drosophila Y chromosome. Mol Biol Evol 31:2612–2623.

Slater GS, Birney E. 2005. Automated generation of heuristics for biological sequence comparison. BMC Bioinformatics 6:31.

RepeatMasker. 2013. Available from: http://www.repeatmasker.org

Smukowski Heil CS, Ellison C, Dubin M, Noor MA. 2015. Recombining without Hotspots: A Comprehensive Evolutionary Portrait of Recombination in Two Closely Related Species of Drosophila. Genome Biol Evol 7:2829–2842.

Song X, Goicoechea JL, Ammiraju JS, Luo M, He R, Lin J, Lee SJ, Sisneros N, Watts T, Kudrna DA, et al. 2011. The 19 genomes of Drosophila: a BAC library resource for genus-wide and genome-scale comparative evolutionary research. Genetics187:1023–1030.

Steinemann M. 1982. Multiple sex chromosomes in Drosophila miranda: a system to study the degeneration of a chromosome. Chromosoma 86:59–76.

Steinemann S, Steinemann M. 2005. Y chromosomes: born to be destroyed. Bioessays 27:10761083.

Sun FL, Cuaycong MH, Craig CA, Wallrath LL, Locke J, Elgin SC. 2000. The fourth chromosome of Drosophila melanogaster: interspersed euchromatic and heterochromatic domains. Proc Natl Acad Sci U S A 97:5340–5345.

Tamura K, Stecher G, Peterson D, Filipski A, Kumar S. 2013. MEGA6: Molecular Evolutionary Genetics Analysis version 6.0. Mol Biol Evol 30:2725–2729.

Thornton K. 2003. Libsequence: a C++ class library for evolutionary genetic analysis. Bioinformatics 19:2325–2327.

Trapnell C, Pachter L, Salzberg SL. 2009. TopHat: discovering splice junctions with RNA-Seq. Bioinformatics 25:1105–1111.

Trapnell C, Roberts A, Goff L, Pertea G, Kim D, Kelley DR, Pimentel H, Salzberg SL, Rinn JL, Pachter L. 2012. Differential gene and transcript expression analysis of RNA-seq experiments with TopHat and Cufflinks. Nat Protoc 7:562–578.

Treangen TJ, Salzberg SL. 2012. Repetitive DNA and next-generation sequencing: computational challenges and solutions. Nature reviews. Genetics 13:36–46.

Urrutia AO, Hurst LD. 2003. The signature of selection mediated by expression on human genes. Genome Res 13:2260–2264.

Vicoso B, Bachtrog D. 2015. Numerous transitions of sex chromosomes in Diptera. PLoS Biol 13:e1002078.

Vicoso B, Bachtrog D. 2013. Reversal of an ancient sex chromosome to an autosome in Drosophila. Nature 499:332–335.

Villasante A, Abad JP, Planello R, Mendez-Lago M, Celniker SE, de Pablos B. 2007. Drosophila telomeric retrotransposons derived from an ancestral element that was recruited to replace telomerase. Genome Res 17:1909–1918.

Wahrman J, Zahavi A. 1955. Cytological Contributions to the Phylogeny and Classification of the Rodent Genus Gerbillus. Nature 175:600–602.

White MJD. 1973. Animal Cytology and Evolution: Cambridge University Press.

Wurster DH, Benirschke K. 1970. Indian Momtjac, Muntiacus muntiak: A Deer with a Low Diploid Chromosome Number. Science 168:1364–1366.

Zhou Q, Bachtrog D. 2012. Sex-specific adaptation drives early sex chromosome evolution in Drosophila. Science 337:341–345.

